# egKnock: identifying direct gene knockout strategies for microbial strain optimization based on metabolic network with gene-protein-reaction relationships

**DOI:** 10.1101/514653

**Authors:** Zixiang Xu

## Abstract

**Background:** Gene knockout method has been used to improve the conversion ratio of industrial strains for many chemical products. There are a series of published algorithms to predict the targets for deletion. Based on metabolic networks, many of these algorithms are designed to predict the target of reaction or enzyme deletion. But as for the many-to-many relationship between genes and reactions, reaction or enzyme deletion is not the ideal strategy for metabolic engineering. GDLS algorithm aims to find direct gene deletion target by using local search, but it actually ignores the logic relationship of gene-protein-reaction.

**Results:** In this study, we aim to find direct gene deletion targets for metabolic network, but the logic relationship of gene-protein-reaction (GPR) is considered. Our algorithm is call egKnock. At the same time, egKnock will provide the solution with multiple strategies and can maximize the minimum target flux of industrial objective in flux variability analysis. We compare egKnock with the algorithm of GDLS and OptORF by predicting the targets of gene deletion for several chemical products with their flux balance analysis testification, flux variability analysis testification and the main flux distribution.

**Conclusions:** By comparison with the algorithm of GDLS and OptORF, we can conclude that egKnock is a nice algorithm for identifying direct gene knockout strategies for microbial strain optimization.

## Background

DNA recombinant and other techniques make it possible to manipulate genetic changes, and gene knockout is one of the methods used to improve the yields of industrial strains for many chemical products. There are a series of published algorithms to predict the targets for deletion [1–6]. Bilevel optimization, which was introduced first time by OptKnock [1], is the core of these algorithms. Based on metabolic networks, many of these algorithms are designed to predict the target of reaction or enzyme deletion, such as OptKnock, ReacKnock [2]. ReacKnock has improved the solving speed and can provide multiple solutions. RobustKnock [3] utilizes triple level optimization method to provide the solution for maximizing the minimum target flux of industrial objective in FVA (flux variability analysis). Of course, as for the many-to-many relationship between gene and reaction, reaction or enzyme deletion is not the ideal strategy for metabolic engineering. GDLS [4] algorithm aims to find direct gene deletion target by using local search, but it actually ignores the logic relationship of gene-protein-reaction (GPR), i.e. it removes all the reactions which a deleted gene concerns. OptORF [5] and OptFlux [6] also aim to find direct gene deletion target, but they are based on metabolic-regulatory integrated network, while this kind of models is actually seldom, and up-to-date only *E.coli* and *Yeast* have the corresponding models [13–15]. Ref [11] reports a modified OptORF without regulatory considerations and it is similar with GDLS in methodology.

In this study, we aim to find direct gene deletion targets for metabolic network, but the logic relationships of gene-protein-reaction (GPR) in metabolic network model are considered. Our algorithm is call egKnock (enzyme gene knockout). At the same time, egKnock will provide the solution with multiple strategies and can maximize the minimum target flux of industrial objective in FVA, while the second function is not included in GDLS, OptFlux and OptORF. The logic relationship of GPR is a tough problem, we firstly transform the model of metabolic network with its GPR relationship to a MILP (mixed integer bilevel linear programming) model by a published Matlab toolbox, named Tiger [7]. Then we utilize an improved bilevel optimization method to make a prediction on the direct gene targets for deletion, while maximizing the minimum target flux of industrial objective in FVA. **Table 1** has shown the comparison among these algorithms about gene deletion prediction.

**Table 1.** Comparison among several algorithms about gene deletion prediction.

| Algorithm | Cell Model | GPR relationship | Maximize min FVA | multiple solutions |
| --- | --- | --- | --- | --- |
| OptKnock | metabolic network | no | no | no |
| ReacKnock | metabolic network | no | no | yes |
| RobustKnock | metabolic network | no | yes | no |
| GDLS | metabolic network | no | no | yes |
| egKnock (this study) | metabolic network | **yes** | **yes** | **yes** |
| OptORF | integrated network | yes | no | yes |
| OptFlux | integrated network | yes | no | yes |

## Methods

### 1) Flux balance analysis and gene-protein-reaction relationship

Flux balance analysis is linear programming (LP) in mathematics, and the objective is usually cell growth, while the constraints are stoichiometric balance constraint and flux boundary constraint. Gene-protein-reaction relationships are logic expressions and it is not convenient to solve a LP with logic expressions as constraints, such as problem (I).

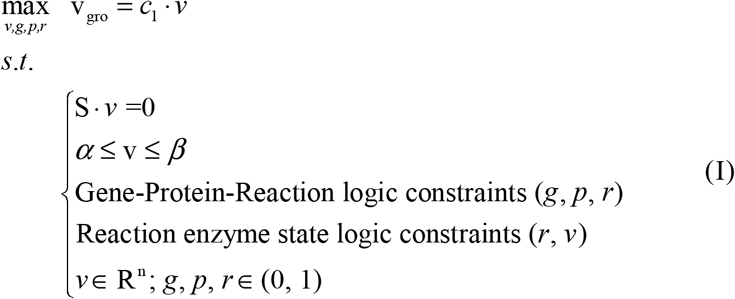

*v* representing fluxes through reactions, *g* representing the Boolean expression state of all genes, *p* prepresenting the presence of each protein, *r* representing the presence of a catalyzing enzyme for each reaction, S is the stoichiometric matrix, α and β represent lower and upper bounds on the fluxes of reaction rates. Reaction enzyme state logic constraints are like: if r_i_=1, α_i_ ≤v_i_ ≤ β_i_; if r_i_=0, v_i_ =0.

### 2) FBA (flux balance analysis) model with GPR

We transform problem (I) where GPRs are logic expressions to the problem (II) where GPRs are inequalities. The logic expressions of GPR relationships include three types “AND, OR, NOT”, and they can be rewritten as linear inequalities [8]. The transforming now can be carried out conveniently by a tool, named Tiger [7].

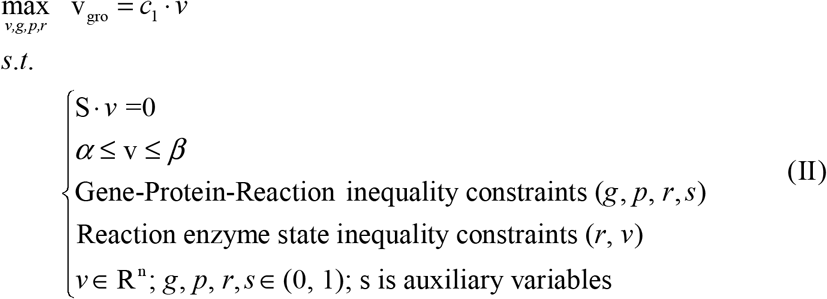

Reaction enzyme state inequality constraints are like: r_i_α_i_ ≤ v_i_ ≤r_i_β_i_.

### 3) Bilevel optimization model with GPR

In order to maximize the rate of product flux, bilevel optimization method can be utilized. The first level is to maximize bioengineering objective, while the second level is to maximize biomass objective. The GPR inequalities and the control constraints are put at the first level. The scale limit of deletion is also in the first level.

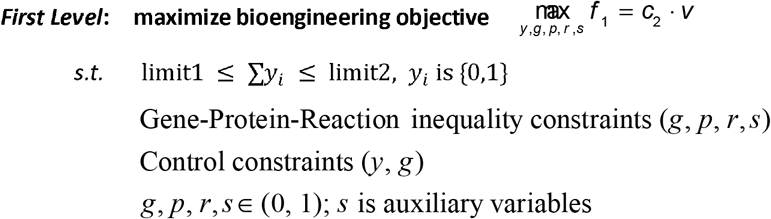

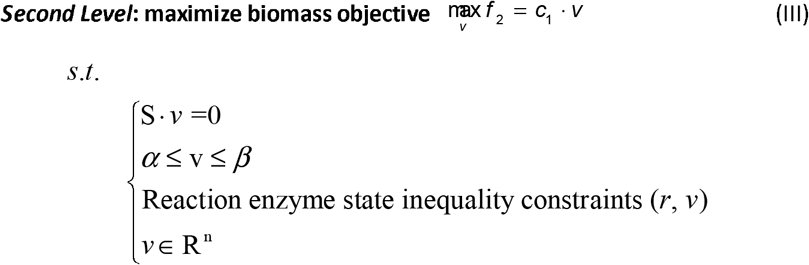

Here, f_1_ is the objective function to maximize industrial production, f_2_ is the objective function for the cell to maximize the growth, i.e. v_gro_; *y* is the control variable, *y*(i)=0 means the gene should be deleted. Control constraints are like: y_i_ = g_i_.

### 4) Maximizing the minimum target flux of industrial objective in FVA

But the gene deletion strategies from the solution of problem (III) only provide the possibility of obtaining a higher yield of product, do not guarantee to maximize the minimum target flux of industrial objective in FVA. In order to make the product flux predicted by egKnock can keep consistent with the minimum target flux in FVA and FBA testification, we change the second level optimization of problem (III) to be minimizing bioengineering objective but under the condition of the same growth, so another set of variables v′ and r’ are introduced. The method we used here is different from RobustKnock [3] which utilizes triple level optimization method.

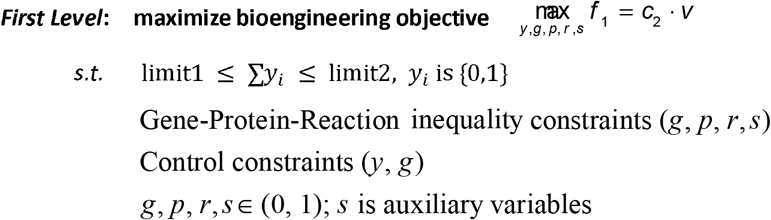

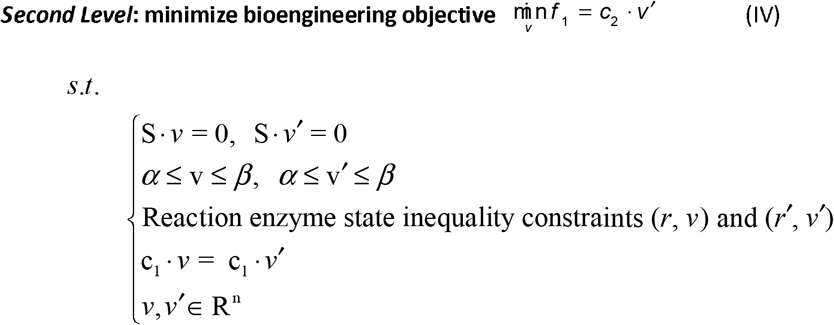

v′ represents fluxes through reactions and it is an equivalent variable of *v*; r’ represents the presence of a catalyzing enzyme for each reaction and it is an equivalent variable of *r*; The objective function of the second level is to minimize bioengineering objective, and the objective function of the first level is to maximize minimized bioengineering objective.

### 5) Transforming bilevel optimization to single level optimization

The above bilevel optimization (IV) can be transformed to a single level MILP (V), and the method utilized is Karush-Kuhn-Tucker (KKT) method, which has been introduced in Ref [2].

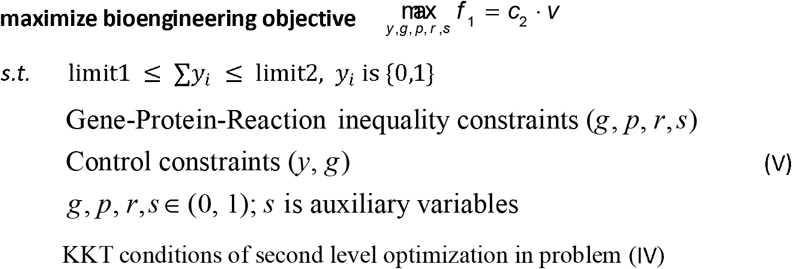

### 6) Multiple solutions

Combinatorial Bender’s cut was used to obtain all the multiple solutions of problem (V), and please refer to Ref [2].

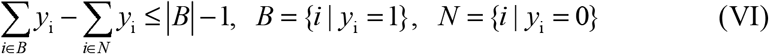

The above MILP (V) together with constraint (VI) can be solved by Gurobi and so on.

## Results and Discussion

In order to evaluate the performance of egKnock with the previous algorithms (GDLS and OptORF), based on a genome-scale metabolic network model of *E. coli*_iAF1260 [9], we applied egKnock to predict gene knockout strategies for producing 5 chemicals (Acetate, Formate, Glycolate, D-Lactate and Fumarate). 15 gene combinatorial deletions were obtained as predicted strategies. Focusing on minimal medium with glucose as sole carbon source, we applied egKnock, GDLS and OptORF respectively towards the production of these different chemicals that can be secreted from *E. coli*. GDLS algorithm in this study was from the corresponding function of COBRAToolbox [10]. But as our experience, it is not convenient to use GDLS for its complex parameter setting and especially it is difficult to reach the given maximum deletion number in the computation. This was also mentioned in Ref [11]. By setting just a constraint of minimum deletion number, we modified the GDLS codes in COBRAToolbox to make it easy to reach the given deletion number. For OptORF, we wrote the codes according to the formula of OptORF paper [11], where it is without regulatory considerations and it is similar with GDLS in methodology. The comparison results under aerobic condition were illustrated in **Table 2**. Here, it was for the reason of computational aspect that we used unified aerobic condition. It is not difficult to get the results under anaerobic condition. Corresponding to **Table 2**, **Table S1** in supplementary information provided the gene names that were deleted and the reactions that were removed according to the GPR relationships. In order to show the capability of multi-solution of egKnock algorithm, **Table 3** has shown the first six alternative solutions for predicting 20-gene deletions to produce Succinate on the *E. coli*_iAF1260. Corresponding to **Table 3**, **Table S2** in supplementary information provided the gene names that were deleted and the reactions that were removed according to the GPR relationships.

**Table 2.**
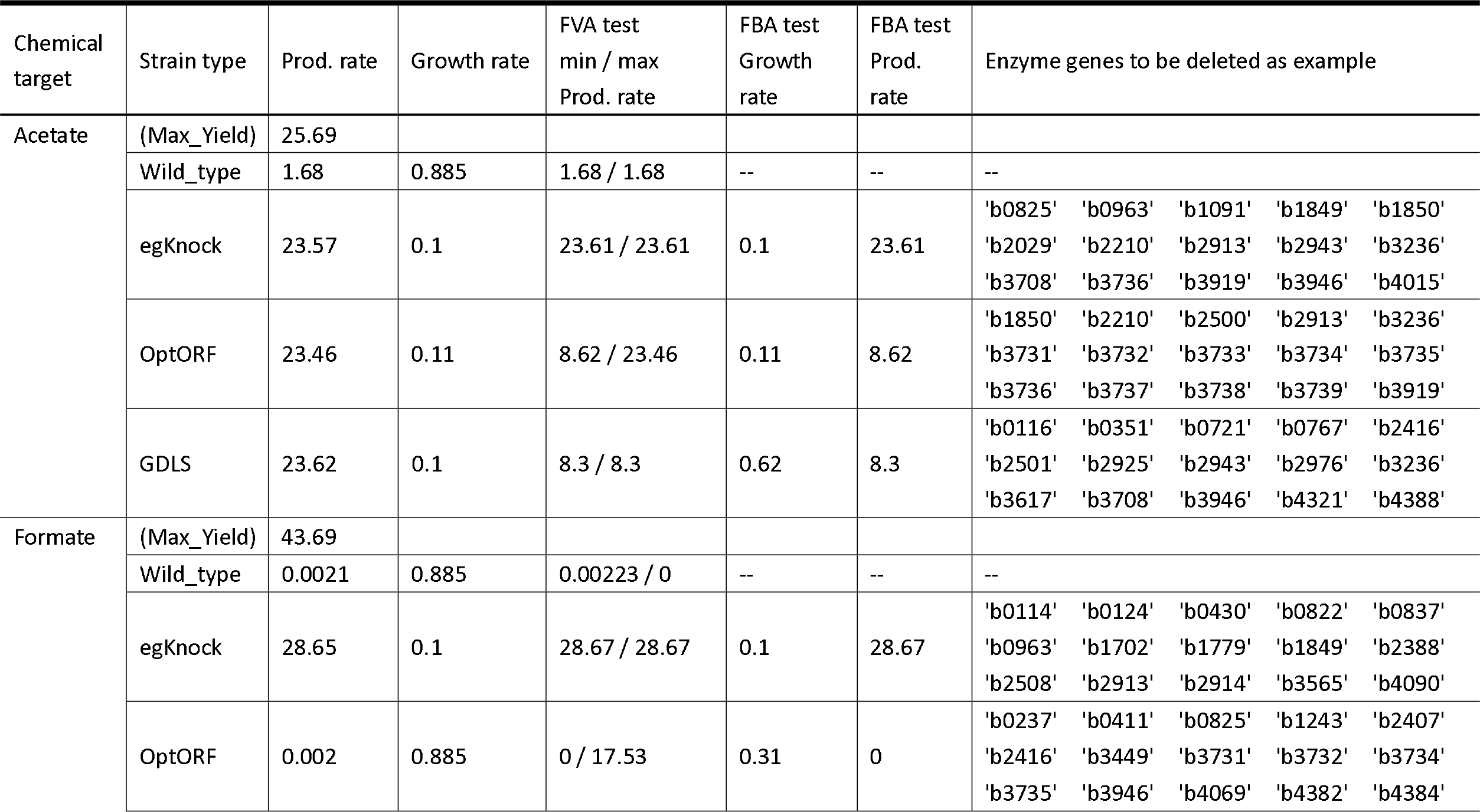
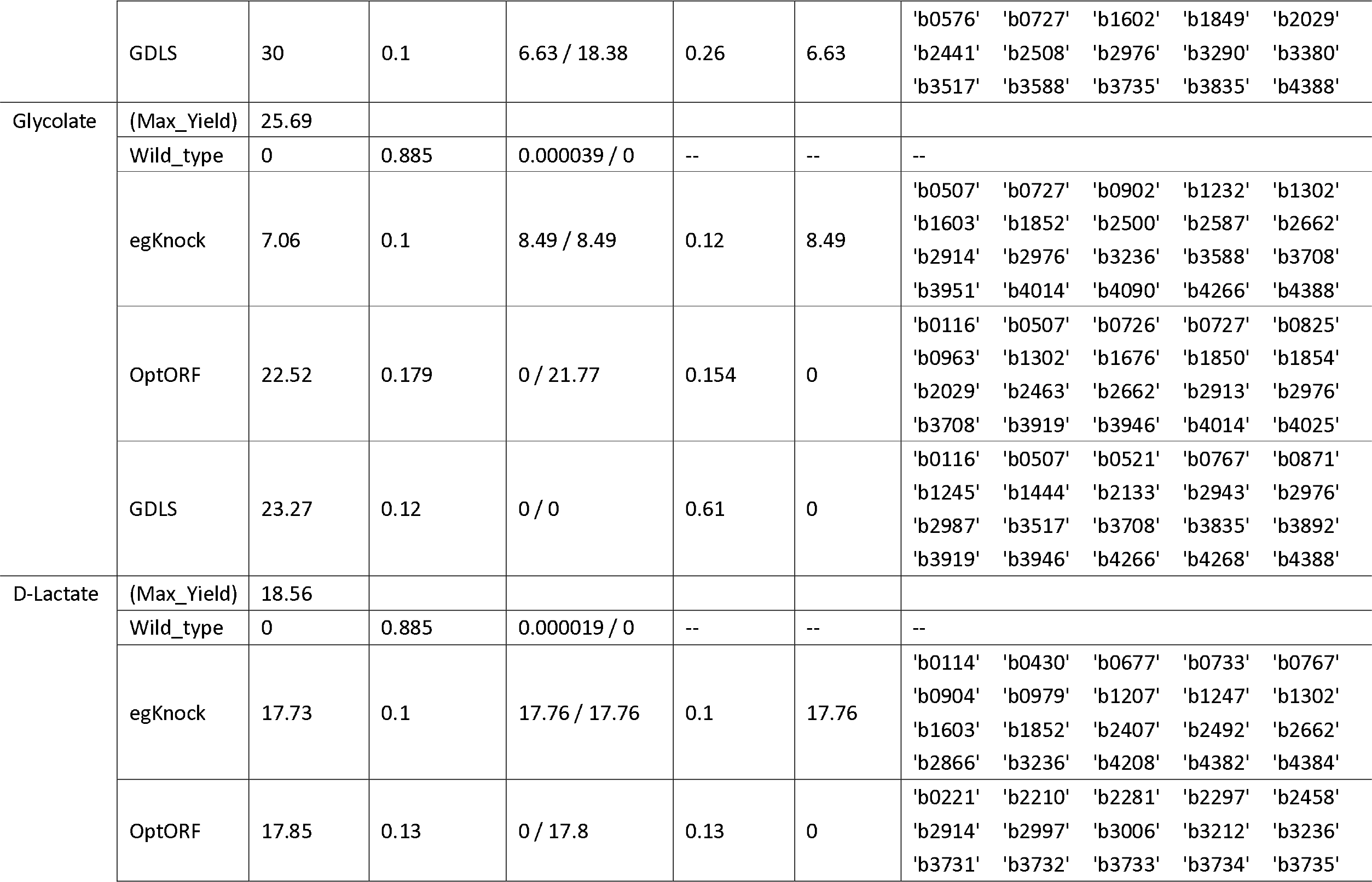
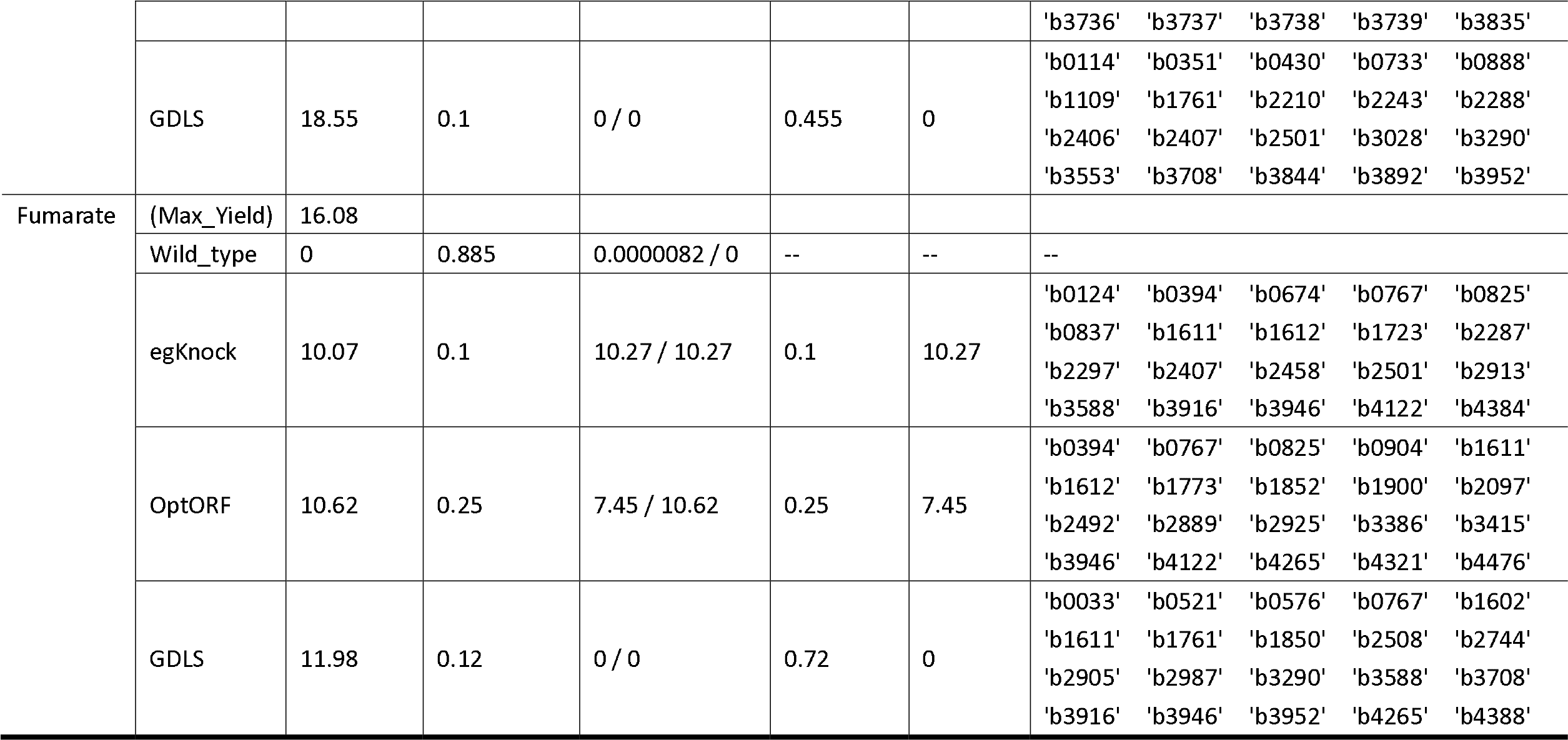
Comparison of the predictions by egKnock, OptORF, GDLS. The following constraints were applied: glucose consumption rate is 10, cell growth is no less than 0.1, maintenance energy metabolism is 8.39, oxygen consumption rate is no higher than 18.5. All the rate unit is mmol/g(Dw)h. Max_yeild means the maximum conversion ratio at the given condition. The maximum computation time was set to 60 min, but the time consumption of most cases are within the set.

**Table 3.** First ten alternative solutions provided by egnock for predicting 20-gene deletions to produce succinate on the model *E. coli*_iAF1260 under aerobic condition with glucose Input = −10 mmol/g(Dw)h. The last two lines are growth rate and product rate respectively.

| No. | 1 | 2 | 3 | 4 | 5 | 6 |
| --- | --- | --- | --- | --- | --- | --- |
| Deletion | 'b0242' | 'b0008' | 'b0124' | 'b0394' | 'b0124' | 'b0124' |
| Strategies | 'b0767' | 'b0112' | 'b0469' | 'b0722' | 'b0323' | 'b0688' |
|  | 'b0904' | 'b0337' | 'b0722' | 'b0767' | 'b0474' | 'b0724' |
|  | 'b1199' | 'b0902' | 'b0767' | 'b0825' | 'b0521' | 'b0751' |
|  | 'b1676' | 'b1199' | 'b0825' | 'b1603' | 'b0529' | 'b0825' |
|  | 'b1761' | 'b1207' | 'b0837' | 'b1723' | 'b0723' | 'b0837' |
|  | 'b1854' | 'b1676' | 'b0910' | 'b1849' | 'b0767' | 'b0910' |
|  | 'b2297' | 'b1761' | 'b1602' | 'b2297' | 'b0825' | 'b1207' |
|  | 'b2407' | 'b1854' | 'b1723' | 'b2388' | 'b0837' | 'b1603' |
|  | 'b2436' | 'b2297' | 'b2297' | 'b2407' | 'b1603' | 'b1723' |
|  | 'b2458' | 'b2407' | 'b2458' | 'b2458' | 'b1723' | 'b1852' |
|  | 'b2492' | 'b2436' | 'b2500' | 'b2501' | 'b2297' | 'b2297' |
|  | 'b2501' | 'b2458' | 'b2661' | 'b2913' | 'b2458' | 'b2458' |
|  | 'b2551' | 'b2464' | 'b2744' | 'b3437' | 'b2501' | 'b2500' |
|  | 'b2987' | 'b2744' | 'b2913' | 'b3588' | 'b2661' | 'b2661' |
|  | 'b3386' | 'b3708' | 'b3588' | 'b3916' | 'b2874' | 'b2690' |
|  | 'b3493' | 'b3738' | 'b3665' | 'b3946' | 'b3588' | 'b3588' |
|  | 'b3736' | 'b3951' | 'b3916' | 'b4265' | 'b3916' | 'b3916' |
|  | 'b4301' | 'b4384' | 'b3946' | 'b4268' | 'b3946' | 'b3946' |
|  | 'b4384' | 'b4388' | 'b4384' | 'b4384' | 'b4388' | 'b4388' |
| growth rate | 0.1 | 0.1 | 0.1 | 0.1 | 0.1 | 0.1 |
| product rate | 9.29 | 9.00 | 11.77 | 11.49 | 11.77 | 11.77 |
| FBA_growth | 0.1 | 0.1 | 0.1 | 0.1 | 0.1 | 0.1 |
| FBA_product | 9.29 | 9.01 | 11.78 | 11.77 | 11.78 | 11.78 |
| FVA_min | 9.29 | 9.00 | 11.78 | 11.77 | 11.78 | 11.78 |
| FVA_max | 9.29 | 9.01 | 11.78 | 11.77 | 11.78 | 11.78 |

Firstly, the product flux predicted by egKnock can keep consistent with the minimum target flux in FVA and FBA testification, while OptORF and GDLS are the worst in this function. When doing the testification of FVA and FBA, we use the standard function “deleteModelGenes” from COBRA Toolbox to remove the predicted target genes and related reactions from the original model. Secondly, egKnock can find direct gene deletion targets for metabolic network, while the logic relationship of GPR is considered. But GDLS actually ignores the logic relationship of GPR, i.e. it removes all the reactions which a gene concerns and this gene is regarded as a deletion target. From the formula (4) of GDLS paper in the Method section, the GPR relationship of metabolic model is reflected in matrix G, but matrix G actually does not include the logic relationships of GPR. That is to say “AND, OR, NOT” logic relationships cannot be reflected in matrix G. OptORF without regulatory considerations [11] is similar to GDLS in methodology. Thirdly, egKnock can return all the alternative deletion strategies in the same search scope with the near industrial objective, while OptORF only provides one deletion strategy for a given deletion number. Fourthly, egKnock adopted numerical method by transforming bilevel optimization to mixed integer program (MIP) and solves MIP by optimization software such as Gurobi [16] or Cplex. Meta-heuristics is another way to solve bilevel optimization [17], where it transforms bilevel optimization to nonlinear program with joint objective and solves it by evolutionary algorithm. OptGene utilizes genetic algorithm to bilevel optimization [18]. Both [17] and [18] are only for metabolic model without considering GRP relationships. In general, numerical methods are faster than heuristics method or genetic algorithms. OptFlux adopts heuristics method as well, so it will spend a long time to reach the optimal point. Fifthly, as we said in Introduction, the model of metabolic-regulatory integrated network is better in describing the behavior of cell than metabolic network model, but this kind of models is actually seldom at present. OptFlux and OptORF are towards integrated network model, but OptORF did not provide the computational tool.

In order to see how the output fluxes are guided to the product after gene deletion, we choose Acetate as an example. We draw respectively the main flux distribution of wild-type E.coli and the main flux distribution of mutant *E.coli* with deleting the 15 genes predicted by egKnock for producing Actate. The flux distribution was calculated with *E.coli*_iAF1260, and when calculating the flux distribution of the mutant, those 15 genes were removed from the model. Of course, the flux distribution of both wild-type *E.coli* and the mutant are complex and we are unable to draw them in one picture clearly at a glance, so we just draw the main flux distribution. The main fluxes mean that the absolute flux values of the reactions in the metabolic network are larger than a given value, such as 5 or 10 mmol/g(Dw)h, then we will draw just several tens of reaction fluxes in one picture. The main flux distribution of wild-type *E.coli* and the mutant are illustrated in **Figure 1** and **Figure 2** respectively, drawn by Paint4Net [12], a COBRA Toolbox extension for visualization of stoichiometric models of metabolism. From the figure, we can see that the flux distribution of mutant *E.coli* is more complex than that of wild-type, but the output fluxes are guided to Acetate. CO2 was the main output flux of wild-type *E.coli* and it was a primary C loss, while this is reduced greatly in the mutant.

**Figure 1.**
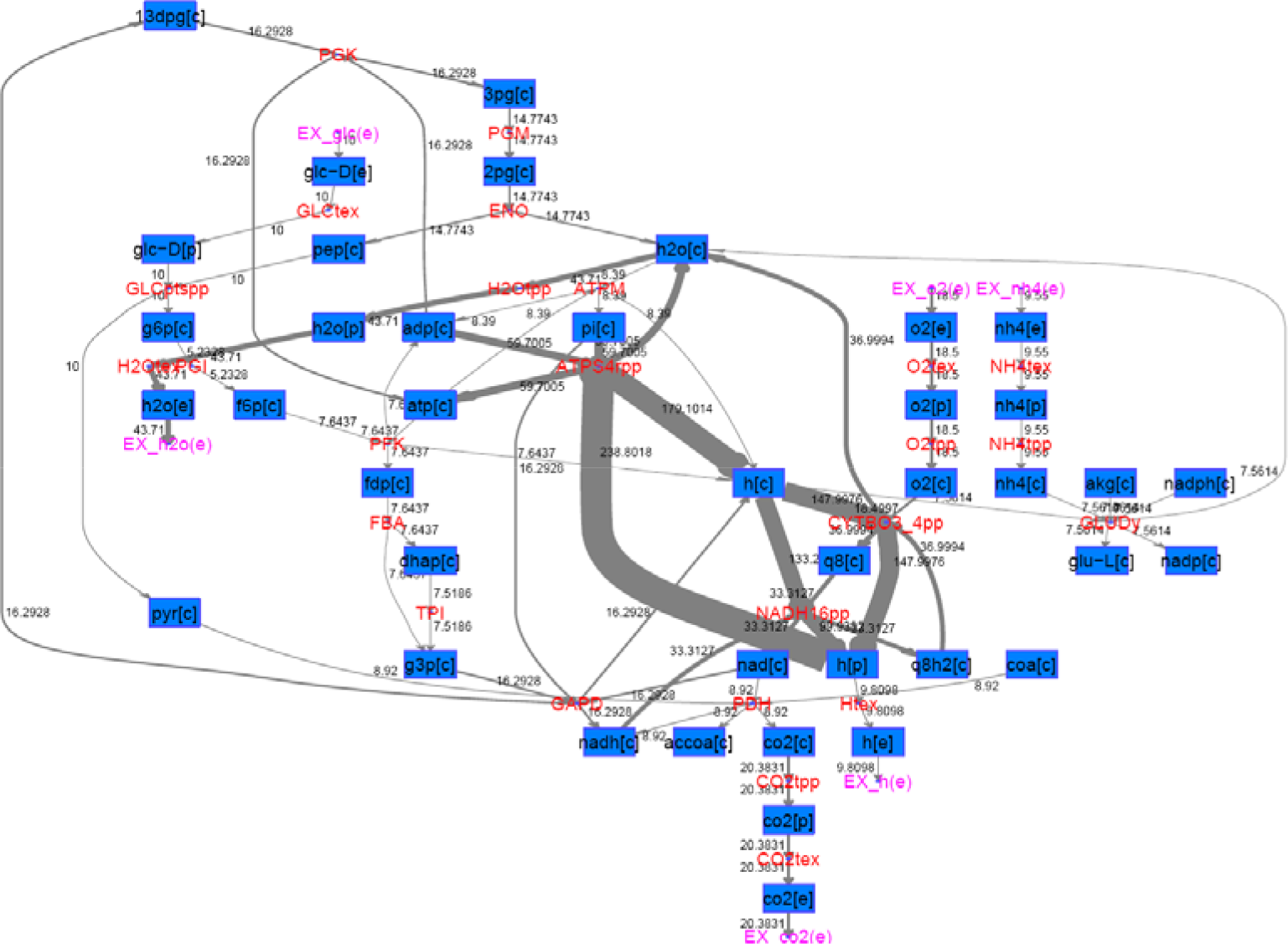
Main flux distribution of wild type *E.coli*. The flux distribution was calculated with *E.coli*_iAF1206 without deleting any genes by setting glucose input as 10, maintenance energy metabolism is 8.39 mmol/g(Dw)h, oxygen consumption rate no higher than 18.5. All the rate unit is mmol/g(Dw)h. Main fluxes mean that the absolute flux values in the reactions are larger than 5. In the figure, blue boxes indicate metabolites, while red circles indicate reactions.

**Figure 2.**
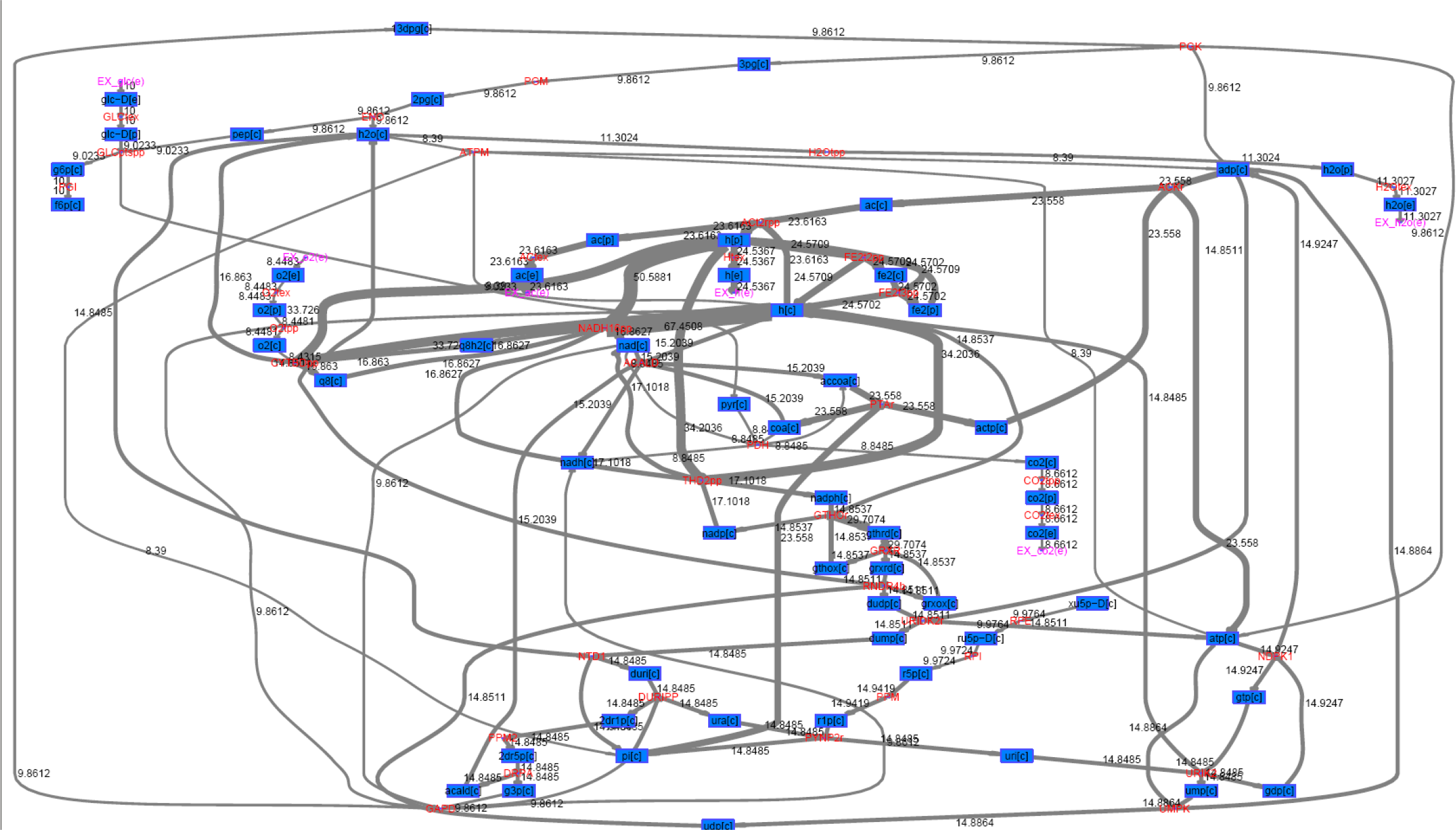
Main flux distribution of mutant *E.coli* with deleting the 15 genes predicted by egKnock for producing Acetate. The flux distribution was calculated with *E.coli*_iAF1206 by setting glucose input as 10, maintenance energy metabolism is 8.39 mmol/g(Dw)h, oxygen consumption rate no higher than 18.5, and at the same time removing those 15 gene from the model. All the rate unit is mmol/g(Dw)h. Main fluxes mean that the absolute flux values in the reactions are larger than 8. In the figure, blue boxes indicate metabolites, while red circles indicate reactions.

## Conclusions

There are several merits of egKnock over GDLS and OptORF. 1) egKnock can guarantee to maximize the minimum target flux of industrial objective in FVA. If a predicted gene deletion strategy which can’t guarantee to maximize the minimum target flux of industrial objective in FVA and this actually just provides the possibility to get a high yield of product, this strategy will actually be an ineffective strategy and we should necessarily design the metabolic pathway in the later. 2) egKnock can find direct gene deletion targets for metabolic network, while the logic relationship of GPR is considered. 3) egKnock can provide all the alternative deletion strategies in the same deletion number and near theoretical yields. It’s very useful for multiple deletion strategies in strain design, for it can provide alternative gene operation strategies. 4) egKnock is stable in running. OptORF is instable in the cases of Formate production, and it can’t provide effective deletion strategies for the chemical productions.

## List of abbreviations

GPR: gene-protein-reaction
FVA: flux variability analysis
egKnock: enzyme gene knockout
MILP: mixed integer bilevel linear programming
linear programming: LP
Karush-Kuhn-Tucker: KKT
FBA: flux balance analysis

## Declarations

### Availability of data and materials

1. The genome-scale metabolic network model of *E. coli*, named iAF1260, could be found in Ref. [9].
2. Supplementary information: including **Table S1** and **Table S2**.
3. The Matlab code of egKnock algorithm, the rewritten Matlab code of GDLS and OptORF algorithms could be obtained by the requirement.

## Authors’ contributions

Conceived and designed the experiments: ZX. Performed the experiments: ZX. Analyzed the data: ZX. Contributed reagents/materials/analysis tools: ZX. Wrote the paper: ZX.

