## Supplemental Tables for "egKnock: identifying direct gene knockout strategies for microbial strain optimization based on metabolic network with gene-protein-reaction relationships"

Table S1. Corresponding to Table 1, the gene names that were deleted and the reactions that were removed according the GPR relationships.

| Chemical target | Strain (mutant) | The names of genes to be deleted | The names of reactions to be removed |
| --- | --- | --- | --- |
| Acetate | egKnock | ' fsaA ' ' mgsA ' ' fabH ' ' purT ' ' eda '  ' gnd ' ' mqo ' ' serA ' ' galP ' ' mdh '  ' tnaA ' ' atpF ' ' tpiA ' ' fsaB ' ' aceA ' | 'ACOATA' 'ATPS4rpp' 'EDA' 'F6PA' 'GALt2pp' 'GART'  'GLCt2pp' 'GND' 'ICL' 'KAS15' 'MDH' 'MDH2' 'MDH3'  'MGSA' 'OAADC' 'PGCD' 'TPI' 'TRPAS2' |
|  | OptORF | ' eda ' ' mqo ' ' purN ' ' serA ' ' mdh '  ' atpC ' ' atpD ' ' atpG ' ' atpA ' ' atpH '  ' atpF ' ' atpE ' ' atpB ' ' atpI ' ' tpiA ' | 'ATPS4rpp' 'EDA' 'GARFT' 'MDH' 'MDH2' 'MDH3'  'OAADC' 'PGCD' 'TPI' |
|  | GDLS | ' lpd ' ' mhpF ' ' sdhC ' ' pgl ' ' ptsI '  ' ppk ' ' fbaA ' ' galP ' ' glcB ' ' mdh '  ' kbl ' ' tnaA ' ' fsaB ' ' gntP ' ' serB ' | 'ACGAptspp' 'ACMANAptspp' 'ACMUMptspp' 'AKGDH'  'ASCBptspp' 'DHAPT' 'FRUURt2rpp' 'FRUpts2pp' 'FRUptspp'  'GALTptspp' 'GALt2pp' 'GAMptspp' 'GLCptspp' 'GLCt2pp'  'GLYAT' 'GLYCL' 'MALTptspp' 'MANGLYCptspp' 'MANptspp'  'MDH' 'MNLptspp' 'PDH' 'PGL' 'PPK2r' 'PPKr'  'PSP_L' 'SBTptspp' 'SUCDi' 'SUCptspp' 'TREptspp'  'TRPAS2' |
| Formate | egKnock | ' aceE ' ' gcd ' ' cyoC ' ' ybiV ' ' yliI '  ' mgsA ' ' pps ' ' gapA ' ' purT ' ' glk '  ' guaB ' ' serA ' ' rpiA ' ' xylA ' ' rpiB ' | 'ALLPI' 'CYTBO3_4pp' 'F6PP' 'G3PT' 'G6PP' 'GAPD' 'GART'  'GLCDpp' 'HEX1' 'IMPD' 'MGSA' 'MN6PP' 'PDH' 'PGCD'  'PPS' 'R5PP' 'RPI' 'XYLI1' 'XYLI2' |
|  | OptORF | ' pepD ' ' tsx ' ' fsaA ' ' oppA ' ' xapA '  ' ptsI ' ' ugpQ ' ' atpC ' ' atpD ' ' atpA '  ' atpH ' ' fsaB ' ' acs ' ' deoA ' ' deoD ' | '4PEPTabcpp' 'ACGAptspp' 'ACMANAptspp' 'ACMUMptspp'  'ACS' 'ADNtex' 'ASCBptspp' 'ATPS4rpp' 'DADNtex'  'DCYTtex' 'DHAPT' 'DURIPP' 'DURItex' 'F6PA' 'FRUpts2pp'  'FRUptspp' 'GALTptspp' 'GAMptspp' 'GLCptspp' 'GPDDA1'  'GPDDA2' 'GPDDA3' 'GPDDA4' 'GPDDA5' 'GUAtex' 'INStex' 'LALGP'  'MALTptspp' 'MANGLYCptspp' 'MANptspp' 'MNLptspp'  'PUNP1' 'PUNP2' 'PUNP3' 'PUNP4' 'PUNP5' 'PUNP6' 'PUNP7'  'SBTptspp' 'SUCptspp' 'TMDPP' 'TREptspp' 'URItex' |
|  | GDLS | ' pheP ' ' sucB ' ' pntB ' ' purT ' ' gnd '  ' eutB ' ' guaB ' ' glcB ' ' trkA ' ' yhfW '  ' gadA ' ' aldB ' ' atpH ' ' ubiB ' ' serB ' | 'AKGDH' 'ALDD2y' 'ALDD3y' 'ATPS4rpp' 'ETHAAL' 'GART' 'GND'  'IMPD' 'OPHHX' 'PSP_L' 'THD2pp' |
| Glycolate | egKnock | ' gcl ' ' sucB ' ' pflA ' ' purU ' ' puuE '  ' pntA ' ' zwf ' ' purN ' ' kgtP ' ' gabT '  ' rpiA ' ' glcB ' ' mdh ' ' aldB ' ' tnaA '  ' pflD ' ' aceB ' ' rpiB ' ' idnO ' ' serB ' | '5DGLCNR' 'ABTA' 'AKGDH' 'AKGt2rpp' 'ALDD2y' 'ALDD3y'  'ALLPI' 'FTHFD' 'G6PDH2r' 'GARFT' 'GLXCL' 'MALS' 'MDH'  'OBTFL' 'PFL' 'PSP_L' 'RPI' 'THD2pp' 'TRPAS2' |
|  | OptORF | ' lpd ' ' gcl ' ' sucA ' ' sucB ' ' fsaA '  ' mgsA ' ' puuE ' ' pykF ' ' eda ' ' pykA '  ' gnd ' ' maeB ' ' gabT ' ' serA ' ' glcB '  ' tnaA ' ' tpiA ' ' fsaB ' ' aceB ' ' pgi ' | 'ABTA' 'AKGDH' 'EDA' 'F6PA' 'GLXCL' 'GLYCL' 'GND'  'MALS' 'ME2' 'MGSA' 'OAADC' 'PDH' 'PGCD' 'PGI' 'PYK'  'TPI' 'TRPAS2' |
|  | GDLS | ' lpd ' ' gcl ' ' ybcF ' ' pgl ' ' poxB '  ' oppC ' ' prr ' ' dld ' ' galP ' ' glcB '  ' pitB ' ' gadA ' ' tnaA ' ' ubiB ' ' fdoI '  ' tpiA ' ' fsaB ' ' idnO ' ' idnK ' ' serB ' | '3PEPTabcpp' '4PEPTabcpp' '5DGLCNR' 'ABUTD' 'AKGDH'  'GALt2pp' 'GLCt2pp' 'GLXCL' 'GLYCL' 'LDH_D2' 'OPHHX' 'PDH'  'PGL' 'POX' 'PSP_L' 'TPI' 'TRPAS2' |
| D-Lactate | egKnock | ' aceE ' ' cyoC ' ' nagA ' ' cydA ' ' pgl '  ' focA ' ' appB ' ' prs ' ' oppF ' ' puuE '  ' pntA ' ' zwf ' ' xapA ' ' focB ' ' gabT '  ' xdhA ' ' mdh ' ' cycA ' ' deoA ' ' deoD ' | '3PEPTabcpp' '4PEPTabcpp' 'ABTA' 'AGDC' 'ALAt2pp' 'BALAt2pp'  'CYTBD2pp' 'CYTBDpp' 'CYTBO3_4pp' 'DALAt2pp' 'DSERt2pp'  'DURIPP' 'FORt2pp' 'FORtppi' 'G6PDH2r' 'HXAND' 'MDH' 'PDH'  'PGL' 'PRPPS' 'PUNP1' 'PUNP2' 'PUNP3' 'PUNP4' 'PUNP5'  'PUNP6' 'PUNP7' 'THD2pp' 'TMDPP' 'XAND' |
|  | OptORF | ' fadE ' ' mqo ' ' nuoI ' ' pta ' ' eutD '  ' rpiA ' ' hybO ' ' exbB ' ' gltB ' ' mdh '  ' atpC ' ' atpD ' ' atpG ' ' atpA ' ' atpH '  ' atpF ' ' atpE ' ' atpB ' ' atpI ' ' ubiB ' | 'ACOAD1f' 'ACOAD2f' 'ACOAD3f' 'ACOAD4f' 'ACOAD5f'  'ACOAD6f' 'ACOAD7f' 'ACOAD8f' 'ADOCBLtonex'  'ATPS4rpp' 'CBItonex' 'CBL1tonex' 'CPGNtonex'  'FE3DCITtonex' 'FE3DHBZStonex' 'FE3HOXtonex' 'FECRMtonex'  'FEENTERtonex' 'FEOXAMtonex' 'GLUSy' 'MDH' 'MDH2'  'MDH3' 'NADH16pp' 'NADH17pp' 'NADH18pp' 'OPHHX' 'PTA2'  'PTAr' |
|  | GDLS | ' aceE ' ' mhpF ' ' cyoC ' ' cydA ' ' trxB '  ' ndh ' ' gdhA ' ' mqo ' ' glpC ' ' nuoA '  ' xapB ' ' xapA ' ' ppk ' ' mdaB ' ' trkA '  ' ghrB ' ' tnaA ' ' fre ' ' fdoI ' ' pflC ' | '2DGLCNRx' '2DGLCNRy' '2DGULRx' '2DGULRy' 'ADNt2rpp'  'CYTBO3_4pp' 'CYTDt2rpp' 'DKGLCNR2x' 'DKGLCNR2y' 'FADRx'  'FE3Ri' 'FLVRx' 'G3PD6' 'G3PD7' 'GLUDy' 'INSt2rpp'  'MDH2' 'MDH3' 'NADH10' 'NADH16pp' 'NADH17pp'  'NADH18pp' 'NADH5' 'NADH9' 'NADPHQR2' 'NADPHQR3'  'NADPHQR4' 'PDH' 'PPK2r' 'PPKr' 'PUNP7' 'THMDt2rpp'  'TRPAS2' 'URIt2rpp' 'XTSNt2rpp' |
| Fumarate | egKnock | ' gcd ' ' mak ' ' asnB ' ' pgl ' ' fsaA '  ' yliI ' ' fumC ' ' fumA ' ' pfkB ' ' nuoB '  ' pta ' ' xapA ' ' eutD ' ' ppk ' ' serA '  ' aldB ' ' pfkA ' ' fsaB ' ' fumB ' ' deoD ' | 'ALDD2y' 'ALDD3y' 'ASNS1' 'F6PA' 'FUM' 'GLCDpp' 'HEX7'  'NADH16pp' 'NADH17pp' 'NADH18pp' 'PFK' 'PFK_2' 'PGCD'  'PGL' 'PPK2r' 'PPKr' 'PTA2' 'PTAr' 'PUNP1' 'PUNP2'  'PUNP3' 'PUNP4' 'PUNP5' 'PUNP6' 'PUNP7' |
|  | OptORF | ' mak ' ' pgl ' ' fsaA ' ' focA ' ' fumC '  ' fumA ' ' ydjI ' ' zwf ' ' araG ' ' fbaB '  ' focB ' ' idi ' ' fbaA ' ' rpe ' ' gntT '  ' fsaB ' ' fumB ' ' idnT ' ' gntP ' ' gntU ' | '5DGLCNt2rpp' 'ARBabcpp' 'F6PA' 'FBA' 'FORt2pp'  'FORtppi' 'FRUURt2rpp' 'FUM' 'G6PDH2r' 'GLCNt2rpp' 'HEX7'  'IDONt2rpp' 'IPDDI' 'PGL' |
|  | GDLS | ' carB ' ' ybcF ' ' pheP ' ' pgl ' ' pntB '  ' fumC ' ' gdhA ' ' eda ' ' guaB ' ' surE '  ' gcvT ' ' pitB ' ' trkA ' ' aldB ' ' tnaA '  ' pfkA ' ' fsaB ' ' pflC ' ' idnT ' ' serB ' | '5DGLCNt2rpp' 'ALDD2y' 'ALDD3y' 'CBPS' 'EDA' 'GLUDy'  'GLYCL' 'IDONt2rpp' 'IMPD' 'NTD10' 'NTD11' 'NTD4' 'NTD7'  'NTD9' 'OAADC' 'PFK_2' 'PGL' 'PSP_L' 'THD2pp' 'TRPAS2' |

Table S2. Corresponding to Table 2, the gene names that were deleted and the reactions that were removed according the GPR relationships.

| No. | 1 | 2 | 3 | 4 | 5 | 6 |
| --- | --- | --- | --- | --- | --- | --- |
| Gene | ' proB ' | ' talB ' | ' gcd ' | ' mak ' | ' gcd ' | ' gcd ' |
| Deletion | ' pgl ' | ' aroP ' | ' apt ' | ' sdhD ' | ' yahI ' | ' pgm ' |
| Strategies | ' focA ' | ' codA ' | ' sdhD ' | ' pgl ' | ' adk ' | ' sdhB ' |
|  | ' dhaL ' | ' pflA ' | ' pgl ' | ' fsaA ' | ' ybcF ' | ' pnuC ' |
|  | ' pykF ' | ' dhaL ' | ' fsaA ' | ' pntA ' | ' folD ' | ' fsaA ' |
|  | ' gdhA ' | ' prs ' | ' yliI ' | ' pfkB ' | ' sdhA ' | ' yliI ' |
|  | ' pykA ' | ' pykF ' | ' cmk ' | ' purT ' | ' pgl ' | ' cmk ' |
|  | ' pta ' | ' gdhA ' | ' pntB ' | ' pta ' | ' fsaA ' | ' prs ' |
|  | ' xapA ' | ' pykA ' | ' pfkB ' | ' glk ' | ' yliI ' | ' pntA ' |
|  | ' hemF ' | ' pta ' | ' pta ' | ' xapA ' | ' pntA ' | ' pfkB ' |
|  | ' eutD ' | ' xapA ' | ' eutD ' | ' eutD ' | ' pfkB ' | ' zwf ' |
|  | ' focB ' | ' hemF ' | ' purN ' | ' ppk ' | ' pta ' | ' pta ' |
|  | ' ppk ' | ' eutD ' | ' gabD ' | ' serA ' | ' eutD ' | ' eutD ' |
|  | ' glyA ' | ' talA ' | ' surE ' | ' gntK ' | ' ppk ' | ' purN ' |
|  | ' pitB ' | ' surE ' | ' serA ' | ' aldB ' | ' gabD ' | ' gabD ' |
|  | ' rpe ' | ' tnaA ' | ' aldB ' | ' pfkA ' | ' yqeA ' | ' yqaB ' |
|  | ' pitA ' | ' atpB ' | ' ade ' | ' fsaB ' | ' aldB ' | ' aldB ' |
|  | ' atpF ' | ' pflD ' | ' pfkA ' | ' idnT ' | ' pfkA ' | ' pfkA ' |
|  | ' sgcE ' | ' deoD ' | ' fsaB ' | ' idnK ' | ' fsaB ' | ' fsaB ' |
|  | ' deoD ' | ' serB ' | ' deoD ' | ' deoD ' | ' serB ' | ' serB ' |
| Reactions | 'ATPS4rpp' | 'ATPS4rpp' | 'ADD' | '5DGLCNt2rpp' | 'ADK1' | 'ALDD2y' |
| to be | 'CPPPGO' | 'CPPPGO' | 'ADPT' | 'ALDD2y' | 'ADK3' | 'ALDD3y' |
| removed | 'DHAPT' | 'CSND' | 'ALDD2y' | 'ALDD3y' | 'ADK4' | 'CYTK1' |
|  | 'FORt2pp' | 'DHAPT' | 'ALDD3y' | 'F6PA' | 'ADNK1' | 'CYTK2' |
|  | 'FORtppi' | 'GLUDy' | 'CYTK1' | 'GART' | 'ALDD2y' | 'F6PA' |
|  | 'GHMT2r' | 'HISt2rpp' | 'CYTK2' | 'GNK' | 'ALDD3y' | 'G6PDH2r' |
|  | 'GLU5K' | 'NTD10' | 'F6PA' | 'HEX1' | 'CBMKr' | 'GARFT' |
|  | 'GLUDy' | 'NTD11' | 'GARFT' | 'HEX7' | 'DADK' | 'GLCDpp' |
|  | 'PGL' | 'NTD4' | 'GLCDpp' | 'IDONt2rpp' | 'F6PA' | 'NMNPtpp' |
|  | 'PIt2rpp' | 'NTD7' | 'NTD10' | 'PFK' | 'GLCDpp' | 'PFK' |
|  | 'PPK2r' | 'NTD9' | 'NTD11' | 'PFK_2' | 'MTHFC' | 'PFK_2' |
|  | 'PPKr' | 'OBTFL' | 'NTD4' | 'PGCD' | 'MTHFD' | 'PGMT' |
|  | 'PTA2' | 'PFL' | 'NTD7' | 'PGL' | 'PFK' | 'PRPPS' |
|  | 'PTAr' | 'PRPPS' | 'NTD9' | 'PPK2r' | 'PFK_2' | 'PSP_L' |
|  | 'PUNP1' | 'PSP_L' | 'PFK' | 'PPKr' | 'PGL' | 'PTA2' |
|  | 'PUNP2' | 'PTA2' | 'PFK_2' | 'PTA2' | 'PPK2r' | 'PTAr' |
|  | 'PUNP3' | 'PTAr' | 'PGCD' | 'PTAr' | 'PPKr' | 'SSALy' |
|  | 'PUNP4' | 'PUNP1' | 'PGL' | 'PUNP1' | 'PSP_L' | 'SUCDi' |
|  | 'PUNP5' | 'PUNP2' | 'PTA2' | 'PUNP2' | 'PTA2' | 'THD2pp' |
|  | 'PUNP6' | 'PUNP3' | 'PTAr' | 'PUNP3' | 'PTAr' |  |
|  | 'PUNP7' | 'PUNP4' | 'PUNP1' | 'PUNP4' | 'SSALy' |  |
|  | 'PYK' | 'PUNP5' | 'PUNP2' | 'PUNP5' | 'SUCDi' |  |
|  | 'RPE' | 'PUNP6' | 'SSALy' | 'PUNP6' | 'THD2pp' |  |
|  |  | 'PUNP7' | 'SUCDi' | 'PUNP7' |  |  |
|  |  | 'PYK' | 'THD2pp' | 'SUCDi' |  |  |
|  |  | 'TALA' |  | 'THD2pp' |  |  |
|  |  | 'TRPAS2' |  |  |  |  |
